# Weight loss improves skeletal muscle mitochondrial energy efficiency

**DOI:** 10.1101/2022.12.21.521461

**Authors:** Patrick J. Ferrara, Marisa J. Lang, Jordan M. Johnson, Shinya Watanabe, Kelsey L. McLaughlin, J. Alan Maschek, Anthony R.P. Verkerke, Piyarat Siripoksup, Amandine Chaix, James E. Cox, Kelsey H. Fisher-Wellman, Katsuhiko Funai

**Affiliations:** Diabetes & Metabolism Research Center, University of Utah; Department of Nutrition & Integrative Physiology, University of Utah; East Carolina Diabetes & Obesity Institute, East Carolina University; Department of Physiology, East Carolina University; Metabolomics Core Research Facility, University of Utah; Molecular Medicine Program, University of Utah; Department of Biochemistry, University of Utah

**Author notes:** Correspondence to: Katsuhiko Funai, Ph.D., Diabetes & Metabolism Research Center, University of Utah.

**Keywords:** Energy efficiency, energy expenditure, mitochondria, oxidative phosphorylation, phospholipids, weight loss

## Abstract

Weight loss is associated with a disproportionate decrease in whole-body energy expenditure that may contribute to the heightened risk for weight-regain. Evidence suggests that this energetic mismatch originates from lean tissue. Although this phenomenon is well documented, the mechanisms have remained elusive. We hypothesized that increased mitochondrial energy efficiency in skeletal muscle is associated with reduced expenditure under weight loss. Wildtype male C57BL6/N mice were fed with high-fat diet for 10 wks, followed by a subset of mice that were maintained on the obesogenic diet (OB) or switched to standard chow to promote weight loss (WL) for additional 6 wks. Mitochondrial energy efficiency was evaluated using high-resolution respirometry and fluorometry. Mass spectrometric analyses were employed to describe the mitochondrial proteome and lipidome. Weight loss promoted ~50% increase in the efficiency of oxidative phosphorylation (ATP produced per O_2_ consumed, or P/O) in skeletal muscle. However, weight loss did not appear to induce significant changes in mitochondrial proteome, nor any changes in respiratory supercomplex formation. Instead, it accelerated the remodeling of mitochondrial cardiolipin (CL) acyl-chains to increase tetralinoleoyl CL (TLCL) content, a species of lipids thought to be functionally critical for the respiratory enzymes. We further show that lowering TLCL by deleting the CL transacylase tafazzin was sufficient to reduce skeletal muscle P/O and protect mice from diet-induced weight gain. These findings implicate skeletal muscle mitochondrial efficiency as a novel mechanism by which weight loss reduces energy expenditure in obesity.

## Introduction

Obesity imposes tremendous health risks but efforts to lose weight is often met with limited success (1, 2). Many find weight loss achieved by weight loss regimens difficult to sustain. As observed in multiple studies including that of the popular “The Biggest Loser” television show (3), it has been postulated that weight loss is associated with an increase in energy efficiency, promoting lower metabolic rate (4–7). These observations are found not only in human weight loss but also in model organisms (8, 9). It is unclear what mechanisms contribute to reduced energy expenditure in the weight-loss state.

Leibel and others demonstrate that weight loss is associated with a decrease in activity-associated energy expenditure in addition to basal metabolic rate (10–13). Skeletal muscle work efficiency appears to be the primary determinant of the disproportionate decline in 24 hr energy expenditure following weight loss, with additional declines in resting energy expenditure (14). Skeletal muscle contributes to the majority of increased energy expenditure induced during physical activity. Previous studies have demonstrated that weight loss may alter skeletal muscle mitochondrial respiration (15–18), though it is unclear if these changes reflect alteration in energy efficiency. In this study we tested our hypothesis that weight loss must improve the efficiency of skeletal muscle mitochondrial respiration.

## Results

### Weight loss reduces whole-body energy expenditure

Wildtype C57BL6/N mice were fed with a Western diet (Envigo, TD.88137) for 10 wks, followed by a subset of mice that were maintained on the obesogenic diet (OB), while others were switched to standard chow to promote weight loss (WL) for an additional 6 wks (Figure 1A). Another group of mice were fed standard chow diet for 16 wks as a lean (LN) reference control group. This strategy successfully and consistently produced groups of mice with divergent body weights and adiposity without altering lean mass (Figure 1B-E).

**Figure 1:**
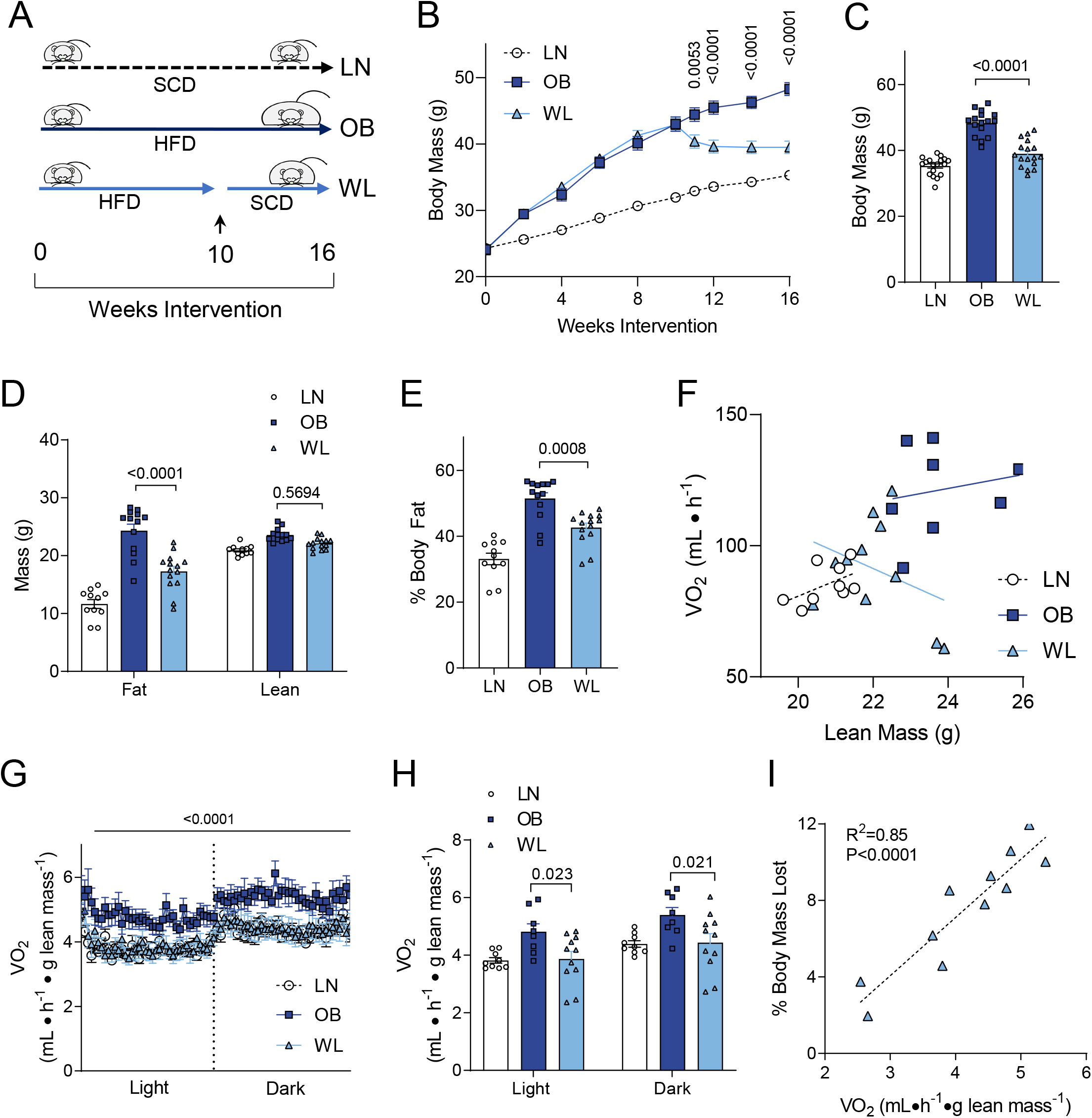
Weight loss promotes a decrease in whole-body energy expenditure. (A) Feeding timeline. LN: mice fed standard chow diet (SCD). OB: mice fed obesogenic high-fat diet (HFD). WL: mice with weight loss induced by switching from HFD to SCD at wk 10. Ad lib fed for all mice. (B) Body mass of LN, OB and WL mice over the 16-week diet intervention. (C) Body mass immediately before terminal experiments. *n*=18 for LN, *n*=16 for OB, *n*=19 for WL for B and C. (D&E) Body composition measured immediately before terminal experiments. *n*=11 for LN, *n*=13 for OB, *n*=14 for WL for both D and E. (F) Relationship between lean mass and total VO_2_ (unnormalized) in LN, OB, and WL groups (*n*=9 for LN, *n*=8 for OB, *n*=11 for WL). (G&H) Whole-body oxygen consumption normalized to lean mass (*n*=9 for LN, *n*=8 for OB, *n*=11 for WL) at wk 15. (I) Pearson correlation analyses of percent body weight loss to the oxygen consumption among the WL group (*n*=11). All data are represented as mean ± SEM. Two-way ANOVA with Sidak multiple comparisons (B,D,G,H) or one-way ANOVA with multiple comparisons (C,E). p-values indicate statistical significance between OB and WL groups.

To assess the energy utilization in these animals, indirect calorimetry experiments were performed at wk 15. Total unnormalized VO_2_ was substantially elevated in the OB group compared to WL or LN group (Figure S1A&B). Correlation analyses indicated that lean mass exhibited a substantially different relationship to total VO_2_ in WL group compared to OB group (Figure 1F, similar relationship with total body mass shown in Figure S1C), suggesting that the lean mass differentially contributes to metabolic rate between OB and WL groups. Indeed, VO_2_ normalized to lean mass were ~20% lower in WL compared to OB group (Figure 1G&H). Spontaneous movement did not explain the differences in VO_2_ (Figure S1D&E). There was no relationship between spontaneous movement and VO_2_ in any of the groups (Figure S1F), suggesting that physical activity does not significantly contribute to 24 hr energy expenditure (19). Respiratory exchange Ratio (RER) was significantly reduced in OB group compared to WL or LN groups (Figure S1G&H), likely representing the lower carbohydrate composition of their diets. Strikingly, among WL mice, there was a very strong positive correlation between VO_2_ to % decrease in body mass (Figure 1G), consistent with the notion that higher energy expenditure contributes to weight loss.

### Weight loss increases skeletal muscle OXPHOS efficiency

Thermogenesis in the brown adipose tissue substantially contributes to whole-body metabolic rate in mice (20). Mitochondrial uncoupling driven by the uncoupling protein 1 (UCP1) resides at the inner mitochondrial membrane to dissipate the proton gradient, driving mitochondrial uncoupling and brown adipose thermogenesis. However, brown adipose tissues from LN, OB, and WL groups did not look different, and had similar mitochondrial density and UCP1 content (Figure S2A).

Previously, we found that the propensity for obesity may be affected by the energy efficiency of sarco/endoplasmic reticulum Ca^2+^-ATPase (SERCA) pump, a highly abundant ATPase that contributes to ~30% of skeletal muscle energy expenditure (21, 22). However, SERCA energy efficiency or abundance in skeletal muscle were not different among the groups (Figure 2A&B). In contrast, weight loss induced a robust increase (~50%) in the efficiency of oxidative phosphorylation (OXPHOS) quantified by ATP produced (energy OUT) per O_2_ consumed (energy IN) or “P/O” in skeletal muscle (Figure 2C). The increase in muscle P/O induced by weight loss was exclusively due to a decrease in O_2_ requirement (Figure S2B) without compromising the capacity for ATP synthesis (Figure S2C). The increase in muscle OXPHOS efficiency was a unique feature of weight loss, as P/O was not different between LN and OB groups.

**Figure 2:**
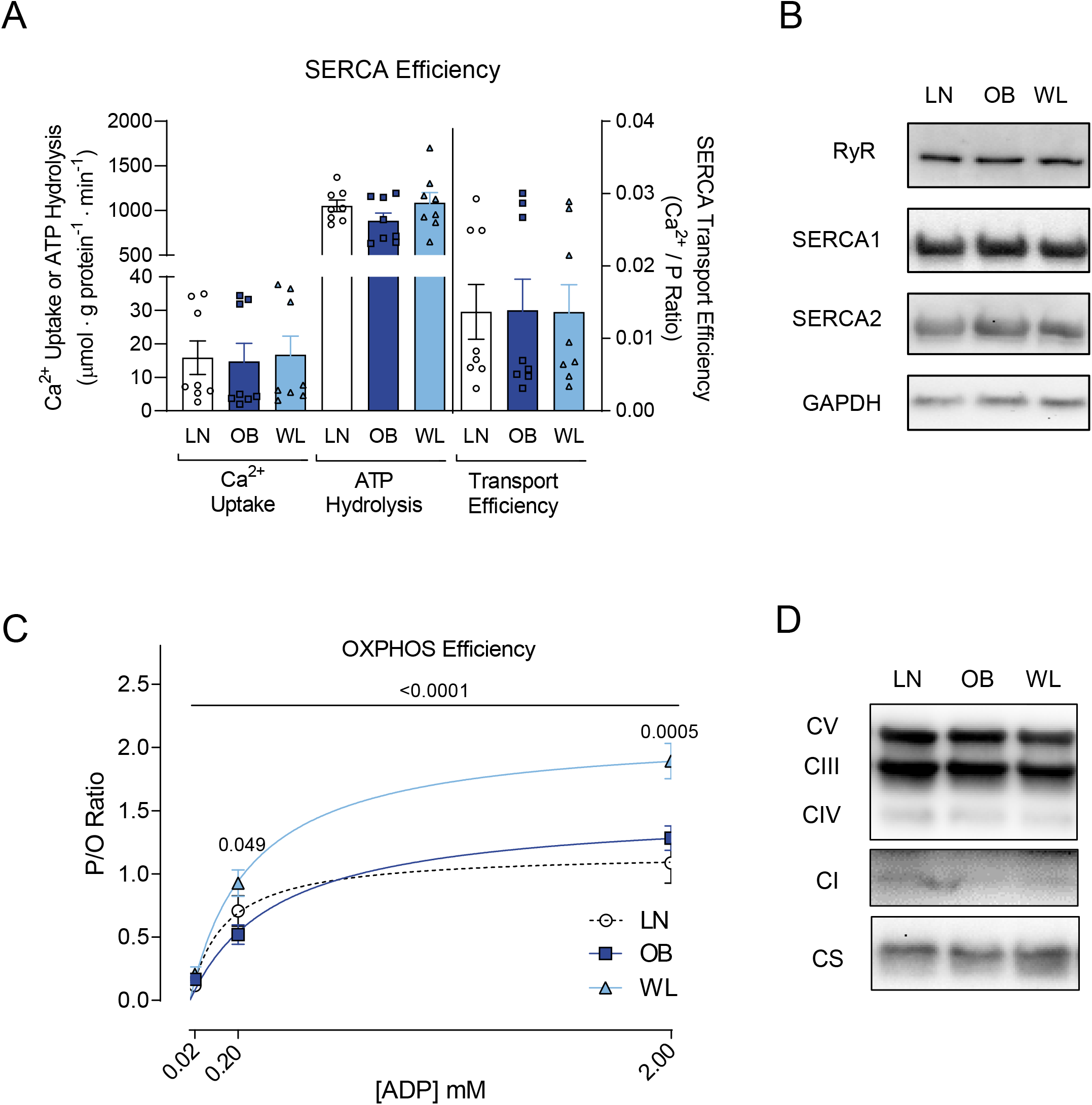
Weight loss improves OXPHOS efficiency in skeletal muscle. (A) Rates of Ca^2+^ uptake, SERCA ATP hydrolysis, and SERCA transport efficiency in skeletal muscle (*n*=8 for all groups). (B) Representative western blots of proteins involved in Ca^2+^ transport. (C) Skeletal muscle P/O ratio in fiber bundles isolated from gastrocnemius muscles (*n*=6 for LN, *n*=8 for OB, *n*=10 for WL). P-values indicate statistical significance between OB and WL groups. (D) Representative western blots of OXPHOS subunits and citrate synthase (CS) in isolated mitochondria. One-way ANOVA with multiple comparisons (A) or two-way ANOVA with Sidak multiple comparisons (C). p-values except in panel I indicate statistical significance between OB and WL groups. p-value in panel I shows statistical significance for correlation. Data are represented as mean ± SEM.

### Weight loss does not alter skeletal muscle mitochondrial proteome

Next, we explored the molecular mechanisms by which weight loss improves skeletal muscle OXPHOS efficiency. Western blot analyses of OXPHOS subunits or citrate synthase did not reveal differences in these proteins in whole tissue lysate (Figure S2D), suggesting these interventions did not alter skeletal muscle mitochondrial content. Western blotting also did not reveal differences in these OXPHOS subunits in isolated mitochondria (Figure 2D). To more comprehensively understand how weight loss influences skeletal muscle OXPHOS enzymes, we analyzed the mitochondrial proteome using mass spectrometry (23). However, weight loss did not induce changes in any of the OXPHOS subunits (Figure 3A-E). In fact, there wasn’t a single mitochondrial protein whose abundance was statistically different between OB and WL group (Figure 3F, comparisons with LN group in Figure S3A&B). Clustering analyses revealed no pattern in the mitochondrial proteome (Figure S3C). We also quantified OXPHOS supercomplex assembly, which also did not reveal any differences (Figure 3G).

**Figure 3:**
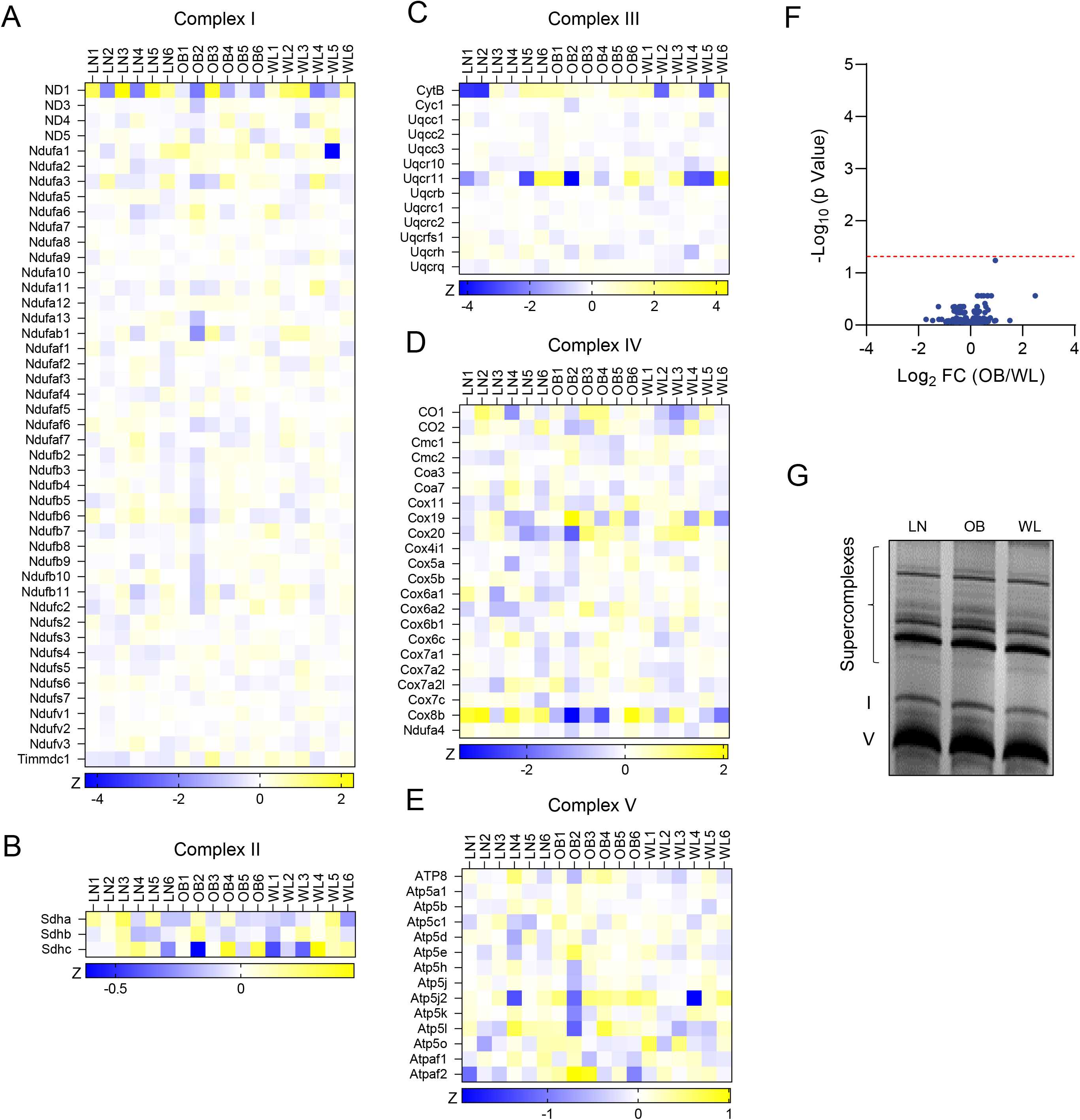
Weight loss does not alter skeletal muscle mitochondrial proteome. (A-E) Heatmap of abundance of OXPHOS subunits measured with mass spectrometry. (F) Volcano plot of differentially abundant mitochondria proteins between OB and WL groups (significance threshold shown with dotted red line). (G) Abundance of respiratory supercomplex formation in isolated mitochondria.

### Weight loss alters skeletal muscle mitochondrial lipidome

OXPHOS enzymes are imbedded in the phospholipid bilayer of the inner mitochondrial membrane (24, 25). Indeed, energy-transducing steps of OXPHOS occurs in (electron transfer) and through (proton transport) the lipid environment. Thus, we examined the mitochondrial lipidome in muscles from LN, OB and WL mice (Figure 4A). Unlike the mitochondrial proteome, there were some changes in the skeletal muscle mitochondrial lipidome induced by weight fluctuation. Importantly, mitochondria from OB and WL groups demonstrated lower lipid-to-protein ratio compared to LN group (Figure 4B), suggesting that muscle mitochondrial membranes become more protein-rich with obesity and remain protein-rich in the weight-loss state. Such change would be predicted to have complex biophysical consequences on OXPHOS dynamics. Unfortunately, it also makes it difficult to interpret differences in mitochondrial lipid composition between LN and the other two groups. For this reason, we decided to focus on the differences in mitochondrial lipidome between the OB and WL groups (Figure S4A-G and Figure 4C). We felt that this is a justifiable strategy given that P/O values were not different between LN and OB groups (Figure 2C), suggesting that differences in mitochondrial membrane lipids between LN and OB is not sufficient to influence OXPHOS efficiency.

**Figure 4:**
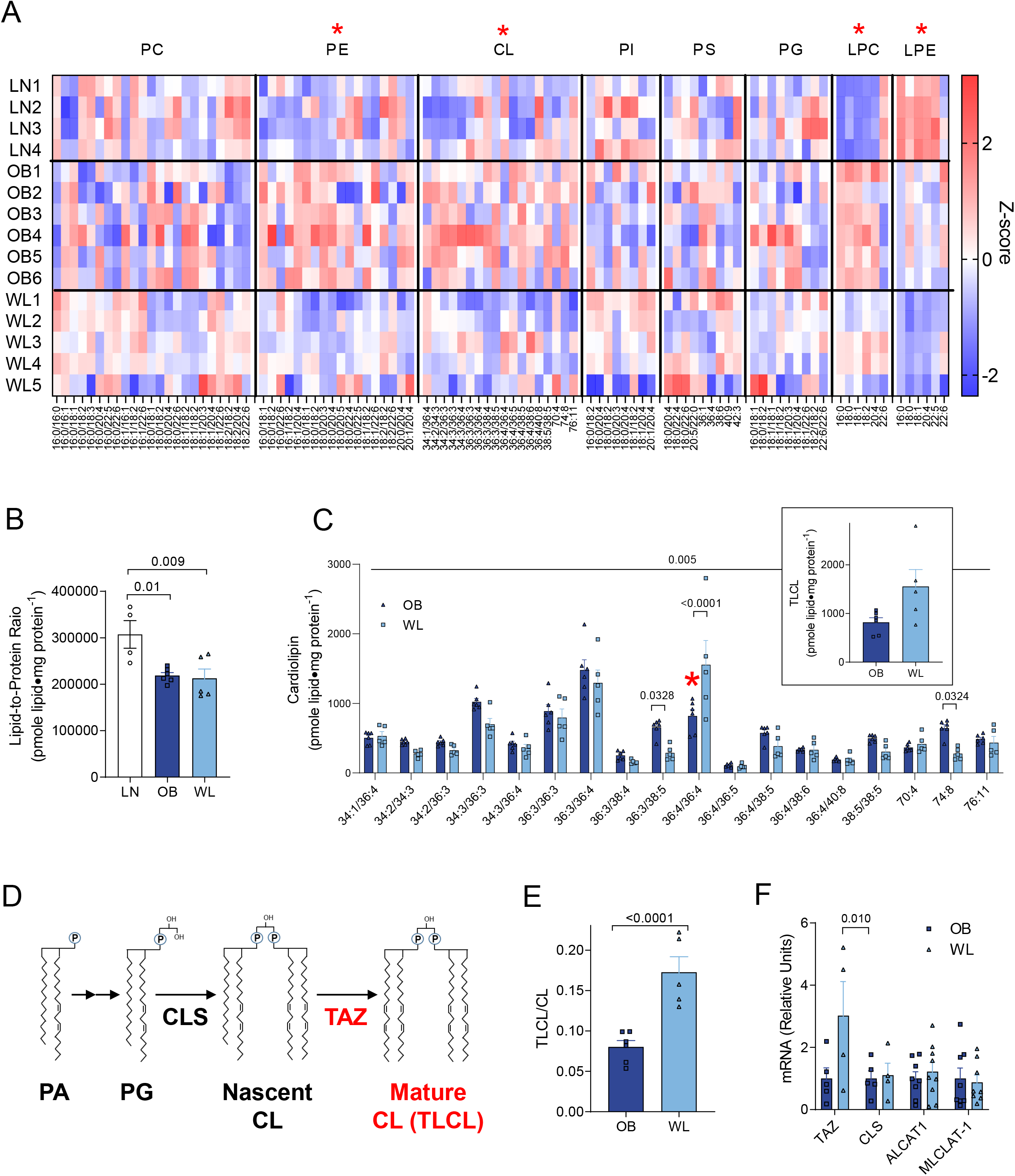
Weight loss accelerates skeletal muscle CL remodeling. (A) Heatmap of relative abundance of mitochondrial lipids. X-axis represents individual lipid species classified according to lipid classes. Red asterisks indicate the main effect of weight loss (OB vs. WL). (B) Lipid-to-protein ratio. *n*=4 for LN, *n*=6 for OB, *n*=5 for WL for A and B. (C) Abundance of individual CL species in isolated mitochondria. TLCL is shown in red asterisk and separately as an insert (*n*=6 for OB, *n*=5 for WL). (D) Schematic of CL synthesis and remodeling that occurs in the inner mitochondrial membrane. (E) TLCL to total CL ratio (*n*=6 for OB, *n*=5 for WL). (F) TAZ, CLS, ALCAT1, and MLCLAT-1 mRNA levels in skeletal muscle (*n*=5 for OB, *n*=4 for WL). One-way ANOVA with multiple comparisons (B), two-way ANOVA with Sidak multiple comparisons (C,F) or unpaired t-test (E). All p-values except for those in panel B indicate statistical significance between OB and WL groups. p-values in panel B show comparisons between LN and OB and LN and WL. Data are represented as mean ± SEM.

Mitochondrial phosphatidylethanolamine (PE) was significantly reduced (Figure 4A and S4B) in WL group compared to the OB group. Mitochondrial PE consists 25-40% of mitochondrial lipids and is primarily synthesized by the phosphatidylserine decarboxylase (PSD) that resides in the inner mitochondrial membrane (26). To determine whether reduction of mitochondrial PE contributes to the increased muscle P/O, we performed lentivirus-mediated knockdown of PSD in murine C2C12 myotubes (Figure S5A). However, PSD knockdown reduced, not increased, mitochondrial P/O (Figure S5B). Thus, reduction in mitochondrial PE observed with weight loss is unlikely to explain the increased P/O.

Weight loss also reduced mitochondrial cardiolipin (CL) in skeletal muscle (Figure 4C). Mitochondrial CL consists 10-20% of mitochondrial lipids and is known to bind with high affinity to OXPHOS enzymes and affect their functions (27). Previous studies suggest that CL may have an impact on OXPHOS efficiency (28–30). CL consists of two phosphatidic acid moieties linked by a central glycerol backbone. CL is almost exclusively localized in the inner mitochondrial membrane, and is synthesized by a series of enzymes localized in the inner mitochondrial membrane including CL synthase (CLS) and CL transacylation enzymes (Figure 4D) (27). For reasons that are not completely understood, CL molecules generated by CLS with uneven acyl-chains are not fully functional and referred as “nascent CL”. These nascent CL molecules are then transacylated to become tetralinoleoyl-CL (TLCL or 18:2/18:2/18:2/18:2-CL), a reaction primarily driven by CL transacylase tafazzin (TAZ) (27, 29). TLCL is thought to be fully functional and referred as “mature CL” (Figure 4D). Thus, we examined the mitochondrial CL portfolio between muscles from OB and WL groups (Figure 4C). Strikingly, even though many of the CL species were lower in WL compared to OB, TLCL was almost twice as highly abundant in WL compared to OB (Figure 4C red asterisk and insert). Indeed, the ratio of TLCL to total CL was 2.2-fold greater in WL compared to OB (Figure 4E). This increase in TLCL content was likely explained by a greater TAZ expression in the WL compared to OB without altering the expression for other CL-synthesizing enzymes (Figure 4F).

### Deficiency in CL remodeling is sufficient to reduce muscle OXPHOS efficiency

We investigated whether TLCL influences OXPHOS efficiency to alter the propensity for weight gain. For these experiments, we utilized mice with doxycycline-induced whole-body knockdown of TAZ (TAZKD mice) compared to doxycycline fed wildtype littermates (Figure 5A). Previous studies have shown that TAZKD mice exhibit greater energy expenditure and are protected from diet-induced obesity (29, 31). Our observations recapitulated these findings (Figure 5B&C). We then examined skeletal muscle tissues from these mice to study the role of CL remodeling in OXPHOS efficiency. As expected, the intervention successfully reduced TAZ expression in skeletal muscle (Figure 5D). We quantified the CL species from skeletal muscle mitochondria using mass spectrometry (Figure 5E). Previous studies showed that virtually all CL species are reduced in TAZKD mice (29, 32), suggesting that nascent CL maybe targeted for degradation without the presence of mature CL. Nevertheless, TAZKD had a disproportionately greater effect to reduce TLCL (Figure 5E, red asterisk) compared to other CL species, shown by a substantial reduction in TLCL to total CL ratio (Figure 5F). The trace amount of TLCL present in these tissues likely arose from alternate CL transacylases such as MLCLAT-1 and ALCAT-1 that are not highly expressed in skeletal muscle. Importantly, we phenotyped skeletal muscle mitochondria from wildtype and TAZKD mice with high-resolution respirometry and fluorometry. Consistent with previous findings, TAZ deletion reduced the capacity for mitochondrial respiration and ATP production (Figure 5G) (29). However, TAZ deletion had a disproportionately greater effect to reduce ATP production than O_2_ consumption, which consequently reduced P/O ratio (Figure 4G). Together, these observations indicate that TAZ deletion is sufficient to reduce skeletal muscle OXPHOS efficiency. In turn, these findings suggest that accelerated CL remodeling in weight loss state may explain the greater OXPHOS efficiency in skeletal muscle.

**Figure 5.**
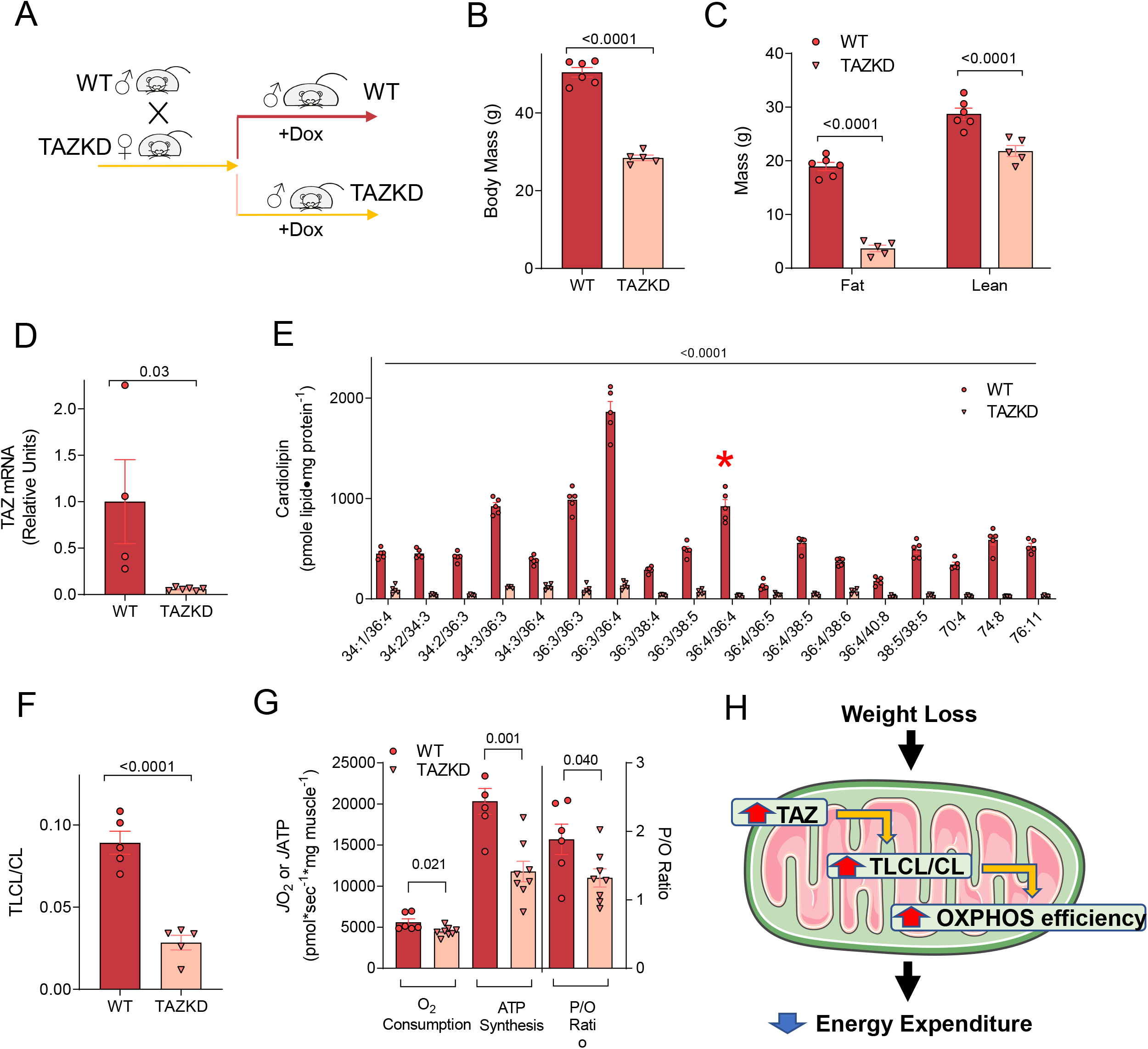
TLCL deficiency reduces OXPHOS efficiency. (A) Schematic of doxycycline intervention in wildtype (WT) and TAZKD littermates. (B) Body mass of WT and TAZKD mice (*n*=6 for WT, *n*=5 for TAZKD). (C) Body composition of WT and TAZKD mice (*n*=6 for WT, *n*=5 for TAZKD). (D) TAZ mRNA abundance in skeletal muscle from WT and TAZKD mice (*n*=4 for WT, *n*=6 for TAZKD). (E) Abundance of individual CL species in isolated mitochondria (*n*=5 for both groups). TLCL is shown in red asterisk. (F) TLCL to total CL ratio (*n*=5 for both groups). (G) Rates for O_2_ consumption, ATP production, and P/O ratio in skeletal muscles from WT and TAZKD mice with 200 μM of ADP (*n*=6 for WT, *n*=8 for TAZKD). (H) Proposed mechanism for how weight loss improves OXPHOS efficiency. Two-way ANOVA with Sidak multiple comparisons (C,E) or unpaired t-test (B,D,F,G). All p-values indicate statistical significance between WT and TAZKD groups. Data are represented as mean ± SEM.

## Discussion

Weight loss reduces whole-body energy expenditure that likely promotes weight regain (4–7). In the current study, we report that weight loss in overweight mice increases skeletal muscle OXPHOS efficiency concomitant to a decrease in whole-body energy expenditure. As one of the organs with large contributions to resting and non-resting energy expenditures, improved skeletal muscle energy efficiency would be predicted to explain a substantial component of a reduction in metabolic rate that occurs with weight loss (33–35). These findings are consistent with previous reports that weight loss is associated with a decrease in activity-associated energy expenditure in addition to basal metabolic rate (10–13).

Skeletal muscle OXPHOS efficiency was increased by ~50% with weight loss. This is a striking increase in energy efficiency that would be predicted to lower muscle energy expenditure, requiring 50% more work to expend equivalent calories. Nevertheless, the increase in OXPHOS efficiency did not coincide with changes in abundance of OXPHOS subunits. Indeed, there was not a single mitochondrial protein whose abundance was significantly affected with weight loss. Instead, weight loss had a more substantial effect on the lipidomic landscape of skeletal muscle mitochondria. One of these changes was an increase in the concentration of mitochondrial TLCL.

We then demonstrated that genetically-induced deficiency in TLCL biosynthesis was sufficient to promote muscle OXPHOS inefficiency and protect mice from diet-induced obesity. These results are consistent with the notion that increased TLCL may contribute to increased OXPHOS efficiency and reduced energy expenditure with weight loss. There are a few caveats to these results pertaining to the use of the TAZKD mice. First, deletion of TAZ is not specific to muscle, so we cannot rule out the possibility that reduced muscle OXPHOS efficiency in these mice could occur indirectly through other tissue. We are currently in a process of developing mice with skeletal muscle-specific knockout of TAZ though these studies are beyond the scope of the current manuscript. Second, doxycycline is known to influence mitochondrial function (36) making this system not ideal for studying bioenergetics. Nevertheless, we treated WT littermates also with doxycycline to control to the best of our abilities. Third, TAZ knockdown lowered the content of all CL species including TLCL, even though TLCL was reduced disproportionately more compared to others. Last but not least, we did not test whether TAZ knockdown would make mice resistant to weight loss-induced reduction in energy expenditure and an increase in muscle OXPHOS efficiency. We plan on performing these experiments as well as the studies on propensity for weight regain in mice with muscle-specific knockout of TAZ.

Findings from this study should not be interpreted to mean that all interventions that promote weight loss increases skeletal muscle OXPHOS efficiency. First, the current study is limited to observations in overweight mice with diet-induced obesity. The results might not be applicable to weight loss in other states. Second, because OB and WL mice were fed diet with different compositions, we cannot rule out the possibility that some of the effects we have observed is not necessarily driven by the effects of weight loss per se. In the WL group, switching to standard chow promptly induced weight loss during the first 2 wks, followed by a 4-wk period of steady body mass. In contrast, OB group continued to gain weight between wks 10 and 16. Third, we did not quantify energy efficiency during muscle contraction, and some evidence suggests a disconnect between P/O and muscle contractile efficiency (37). Last but not least, the current study was performed in room temperature. It would be important to study how these findings are recapitulated in thermoneutrality. It is important to interpret our findings in the context of these caveats.

In summary, weight loss promotes an increase in skeletal muscle OXPHOS efficiency that likely contributes to reduced whole-body energy expenditure. Weight loss also coincided with increased mitochondrial TLCL, and deficiency of TLCL was sufficient to reduce muscle OXPHOS efficiency and make mice more resistant to weight gain. We interpret these findings to propose that weight loss accelerates CL remodeling in skeletal muscle to improve OXPHOS efficiency and lower whole-body energy expenditure (Figure 5H). We speculate that such decrease in metabolic rate arose from a tremendous evolutionary pressure to conserve energy in states of energy deprivation. In our age, these adaptive responses likely strongly contribute to increased propensity for a rebound in adiposity.

## Materials and methods

### Animals and Diet Intervention

Male C57BL/6NCrl (Charles River: 027) mice were used for the weight loss study. At 10 weeks of age mice were either maintained on standard chow diet (SCD; Envigo 2920X) or fed high fat diet (42% calories from fat; HFD; Envigo: TD88137). After 10 weeks of HFD feeding a subset of mice were switched back to SCD while the others continued HFD feeding for another 6 weeks. Heterozygous TAZKD mice were obtained from the Jackson Laboratory (Stock number 014648). TAZ knockdown was induced *in utero* by supplying 625 mg/kg doxycycline chow (Envigo, TD.09628) as previously described (38). Briefly, female TAZKD mice were maintained on doxycycline chow (625 mg/kg) at least 5 days before being mated with male wildtype mice. The doxycycline diet was removed during the mating period, and following copulation, males were removed and the doxycycline diet was reintroduced for the duration of gestation. Following weaning all offspring were maintained on the doxycycline diet for the remainder of the study. For all experiments, mice were provided access to food ad libitum, maintained on a 12-hour light/dark cycle, and fasted for ~4 hours prior to terminal experiments. For terminal experiments mice were given intraperitoneal injection of 80 mg/kg ketamine and 10 mg/kg xylazine, after which tissues were harvested. All animals were randomized and no animals were excluded from the analyses. All procedures were approved by the University of Utah Institutional Animal Care and Use Committee.

### Metabolic Cage & Body Composition

Whole mouse indirect calorimetry and body composition were measured as previously described (21). Columbus Instruments Lab Monitoring Systems were used to measure VO_2_, VCO_2_, respiratory exchange ratio (RER; VCO_2_/VO_2_), and activity. Mice were housed individually and acclimated for at least 24 hr before data collections. Data from the final complete light/dark cycle were used for analysis. Bruker Minispec NMR (Bruker, Germany) was used to determine composition of fat and fat-free mass.

### Mitochondrial and Sarco/endoplasmic Reticulum Enrichment

Gastrocnemius muscles were used to isolate fractions enriched in mitochondria or sarco/endoplasmic reticulum (SR) as previously described (21, 29). For mitochondrial enrichment, muscles were minced in mitochondrial isolation medium (300 mM sucrose, 10 mM HEPES, 1 mM EGTA, and 1 mg/mL BSA) and subsequently homogenized using a Teflon-glass system. Homogenates were then centrifuged at 800 x g for 10 min, after which the supernatant was taken and centrifuged at 12,000 x g for 10 min. The resulting mitochondrial pellet was carefully resuspended in mitochondrial isolation medium without BSA. For SR isolations, muscles were homogenized [300 mM sucrose, 20 mM HEPES pH 7.4, Halt protease (78430)] and underwent differential centrifugation (1,300 x g for 10 min, 20,000 x g for 20 min, 180,000 x g for 2 hr 15 min) (39) to pellet an SR-enriched fraction, which was resuspended in SR isolation buffer.

### High-Resolution Respirometry and Fluorimetry

Respiration in permeabilized muscle fiber bundles was performed as previously described (26, 29). Briefly, a small portion of freshly dissected red gastrocnemius muscle tissue was placed in buffer X [7.23 mM K_2_EGTA, 2.77 mM Ca K_2_EGTA, 20 mM imidazole, 20 mM taurine, 5.7 mM ATP, 14.3 mM phosphocreatine, 6.56 mM MgCl_2_.6H_2_O, and 50 mM K-MES (pH 7.1)]. Fiber bundles were separated and permeabilized for 30 min at 4°C with saponin (30 μg/mL) and immediately washed in buffer Z [105 mM K-MES, 30 mM KCl, 10 mM K_2_HPO_4_, 5 mM MgCl_2_ 6H_2_O, BSA (0.5 mg/mL), and 1 mM EGTA (pH 7.4)] for 15 min. After washing, high-resolution respiration rates were measured using an OROBOROS Oxygraph-2k. The muscle fibers were suspended in buffer Z with 20 mM creatine monohydrate and 10 μM blebbistatin to inhibit myosin adenosine triphosphatases during respiration measurements. Fiber bundles were added to the oxygraph chambers containing assay buffer (105 mM MES potassium salt, 30 mM KCl, 10 mM K_2_HPO_4_, 5 mM MgCl_2_, 0.5 mg/mL BSA). Respiration was measured in response to the following substrates: 0.5 mM malate, 5 mM pyruvate, 5 mM glutamate, 10 mM succinate, 1.5 μM FCCP. ATP production was measured fluorometrically using a Horiba Fluoromax-4 (Horiba Scientific), by enzymatically coupling ATP production to NADPH synthesis as previously described (40). Respiration and ATP production were measured in the presence of 20, 200, and 2000 μM ADP.

### Sarco/Endoplasmic Reticulum ATPase Efficiency Assay

Sarco/endoplasmic reticulum ATPase (SERCA) efficiency assays were performed as previously described (21). SR-fraction was quantified using BCA protein assay (Pierce, 23225) and 10 μg of SR protein was used in each replicate for SERCA-dependent Ca^2+^-uptake and ATPase activity assay. SERCA-dependent Ca^2+^ uptake and ATPase activity assays were performed in buffer containing 60 mM HEPES, 200 mM KCl, 15 mM MgCl_2_, 10 mM NaN_3_, 1 mM EGTA, and 0.005% Triton-X at a pCa of 5.15 and with or without 15 μM thapsigargin. Free Ca^2+^ was determined using Maxchelator Ca-Mg-ATP-EDTA Calculator using constants from the NIST database #46 v8 at 37°C, pH 7.3, and ionic constant of 0.25N (41). ATP SERCA-dependent measures were calculated by taking the difference of values without thapsigargin to values with thapsigargin [Total (without thapsigargin) – SERCA independent (with thapsigargin) = SERCA-dependent]. Ca^2+^ uptake assay buffer additionally contained 5 mM of (COOK)_2_. ATPase activity assay buffer additionally contained 10 mM PEP, 1.5 mM NADH, 2.4–4 units of pyruvate kinase/ 3.6–5.6 units lactate dehydrogenases enzymes (Sigma, P0294).

Calcium uptake assays were performed as previously described (22). Reactions were started by the addition of 4 mM ATP and ^45^CaCl_2_ (Perkin Elmer, NEZ013001MC) to assay buffer with sample. After incubation for 15 minutes at 37°C with 300 RPM rotation, assay was quenched with the addition of 150 mM KCl and 1 mM LaCl_3_ and placed on ice. Samples were then filtered on to a 0.22 μm PES membrane filter (Millipore, GPWP02500), rinsed 3 × 5 mL PBS, and processed for scintillation counting. SERCA ATPase activity assay (42) was performed on a 96-well plate reader. The assay was initiated by the addition of 4 mM ATP and the absorbance at 340 nm was recorded every 60 seconds for 30 minutes at 37°C. SERCA transport efficiency was determined by the ratio of SERCA-dependent Ca^2+^ uptake to SERCA-dependent ATPase hydrolysis.

### Western Blot

Western blots were performed as previously described (43). Protein homogenates were analyzed for abundance of ryanodine receptor (RyR; Santa Cruz 13942), Sarco/Endoplasmic Reticulum Calcium ATPase 1 (SERCA1; Abcam 2818), SERCA2 (Abcam 3625), Glyceraldehyde 3-phosphate dehydrogenase (GAPDH: Cell Signal 2118), mitochondrial complexes I-V (Abcam 110413), citrate synthase (Abcam 96600), and uncoupling protein-1 (UCP1: Alpha Diagnostic UCP11-A).

### Sample Preparation and nLC-MS/MS Label Free Proteomic Analysis

Mitochondria were purified and subjected to label free proteomic screening as previously described (44). Isolated mitochondria were lysed in Buffer D (8 M urea in 40 mM Tris, 30 mM NaCl, 1 mM CaCl_2_, 1 × cOmplete ULTRA mini EDTA-free protease inhibitor tablet; pH = 8.0), as described previously.(23) The samples were subjected to three freeze–thaw cycles, and sonication with a probe sonicator in three 5 s bursts (Q Sonica #CL-188; amplitude of 30). Samples were then centrifuged at 10,000 × g for 10 min at 4°C. Protein concentration was determined by BCA protein assay. Equal amounts of protein were reduced with 5 mM DTT at 37°C for 30 min, and then alkylated with 15 mM iodoacetamide at room temperature for 30 min in the dark. Unreacted iodoacetamide was quenched with DTT up to 15 mM. Initial digestion was performed with Lys C (ThermoFisher Cat# 90,307; 1:100 w:w; 2 μg enzyme per 200 μg protein) for 4 h at 37°C. Following dilution to 1.5 M urea with 40 mM Tris (pH = 8.0), 30 mM NaCl, 1 mM CaCl_2_, samples were digested overnight with trypsin (Promega; Cat# V5113; 50:1 w/w, protein:enzyme)at 37°C. Samples were acidified to 0.5% TFA and then centrifuged at 4000×*g* for 10 min at 4°C. Supernatant containing soluble peptides was desalted, as described previously (23) and then eluate was frozen and lyophilized.

Final peptides were suspended in 0.1% formic acid, quantified (ThermoFisher Cat# 23,275), and then diluted to a final concentration of 0.25 μg/μL. Samples were subjected to nLC-MS/MS analysis using an UltiMate 3000 RSLCnano system (ThermoFisher) coupled to a Q Exactive Plus Hybrid Quadrupole-Orbitrap mass spectrometer (ThermoFisher) via a nanoelectrospray ionization source. For each injection, 4 μL (1 μg) of sample was first trapped on an Acclaim PepMap 100 20 mm × 0.075 mm trapping column (ThermoFisher Cat# 164,535; 5 μL/min at 98/2 v/v water/acetonitrile with 0.1% formic acid). Analytical separation was then performed over a 95 min gradient (flow rate of 250nL/min) of 4–25% acetonitrile using a 2 μm EASY-Spray PepMap RSLC C18 75 μm × 250 mm column (ThermoFisher Cat# ES802A) with a column temperature of 45°C. MS1 was performed at 70,000 resolution, with an AGC target of 3 × 106 ions and a maximum injection time (IT) of 100 ms. MS2 spectra were collected by data-dependent acquisition (DDA) of the top 15 most abundant precursor ions with a charge greater than 1 per MS1 scan, with dynamic exclusion enabled for 20 s. Precursor ions isolation window was 1.5 m/z and normalized collision energy was 27. MS2 scans were performed at 17,500 resolution, maximum IT of 50 ms, and AGC target of 1 × 105 ions.

Proteome Discoverer 2.2 (PDv2.2) was used for raw data analysis, with default search parameters including oxidation (15.995 Da on M) as a variable modification and carbamidomethyl (57.021 Da on C) as a fixed modification. Data were searched against the Uniprot Mus musculus reference proteome (Proteome ID: UP 000000589), as well as the mouse Mito Carta 2.0 database (45). PSMs were filtered to a 1% FDR and grouped to unique peptides while maintaining a 1% FDR at the peptide level. Peptides were grouped to proteins using the rules of strict parsimony and proteins were filtered to 1% FDR. Peptide quantification was done using the MS1 precursor intensity. Imputation was performed via low abundance resampling. Using only high confidence master proteins, mitochondrial enrichment factor (MEF) was determined by comparing mitochondrial protein abundance (i.e., proteins identified to be mitochondrial by cross-reference with the MitoCarta 2.0 database) to total protein abundance.

### Mitochondrial Lipid Mass Spectrometry

Lipids were extracted from mitochondrial enriched fractions as previously described (26) with internal standards (Avanti Polar Lipids: 330707). Untargeted mass spectrometry was performed (Agilent 6530 UHPLC-QTOF mass spectrometer) and analyzed in negative (lyso-PC, lyso-PE, PC, PE, PS, PI, and PG) or positive (CL) modes. Lipid content reported is normalized to mitochondrial protein content.

### Native PAGE

Mitochondrial supercomplex analysis was performed as previously described (26). Isolated mitochondria (100 μg) suspended in MIM were pelleted at 12,000 x g for 15 min and subsequently solubilized in 20 μL sample buffer (4% digitonin, 1x native PAGE sample buffer) for 20 min on ice and then centrifuged at 20,000 x g for 30 min at 4°C. 15 μL of the supernatant (75 μg) was collected and placed into a new tube and mixed with 2 μL G-250 sample buffer additive. The samples and standards were then loaded onto a native PAGE 3 to 12% Bis-Tris Gel (BN1001BOX, Thermo Fisher Scientific), and electrophoresis was performed at 150 V for 3 hours on ice. The gel was then placed in de-stain solution (20% methanol and 10% acetic acid) then shaken on an orbital shaker for 10 min at room temperature. De-stain solution was discarded and fresh de-stain solution was added before shaking for 60 min at room temperature. After incubation, the gel was placed in de-staining solution and incubated overnight at 4°C on an orbital shaker.

### Cell Culture

C2C12 myoblasts were grown [high glucose DMEM + 10% fetal bovine serum (FBS) + 100 μg/ml of penicillin/streptomycin] and differentiated into myotubes [low glucose DMEM (1 g/L glucose, L-glutamine, 110 mg/L sodium pyruvate) + 2% horse serum + 100 μg/ml of penicillin/streptomycin]. HEK 293T cells were maintained in high glucose DMEM + 10% FBS + 100 μg/ml of penicillin/streptomycin. Lentivirus-mediated knockdown of PSD was performed as previously described (26). Vectors were sourced from Sigma (St. Louis, MO) for shRNA for mouse PISD (shPSD: TRCN0000115415), and Addgene (Cambridge, MA) for psPAX2 (ID #12260), pMD2.G (ID #12259), and scrambled shRNA plasmid (SC: ID #1864).

### Quantitative PCR

Samples were homogenized in TRIzol reagent (Life Technologies) to extract total RNA. One microgram RNA was reverse-transcribed using an IScript cDNA synthesis kit (Bio-Rad). Reverse transcription PCR (RT-PCR) was performed with the Viia 7 Real-Time PCR System (Life Technologies) using SYBR Green reagent (Life Technologies). All data were normalized to ribosomal L32 gene expression and were normalized to the mean of the control group. Primers were based on sequences in public databases.

### Statistics

Statistical analysis was performed using GraphPad Prism 9 software. One-way ANOVA with multiple comparisons, two-way ANOVA with Sidak multiple comparisons, Pearson correlation analyses, or unpaired t-test were performed for group comparisons. Where multiple comparisons were made, p-values between OB and WL groups or WT and TAZKD groups are shown in the figures (see legends). Simple linear regressions were performed for correlation analysis. All data are represented as mean ± SEM and statistical significance was set at P ≤ 0.05.

## Acknowledgments

Author contributions: P.J.F, M.J.L, and K.F. designed the study. P.J.F., M.J.L., J.M.J., and S.W. performed the mouse studies and performed mitochondrial phenotyping experiments. A.R.P.V. performed the SERCA experiments. K.L.M and K.H.F-W performed the mitochondrial proteomic analyses. J.A.M. and J.E.C. performed the mitochondrial lipidomic analyses. P.S. performed the cell culture studies. P.J.F. and K.F. wrote the manuscript with edits from all authors.

---

**Supplemental Figure S1.**
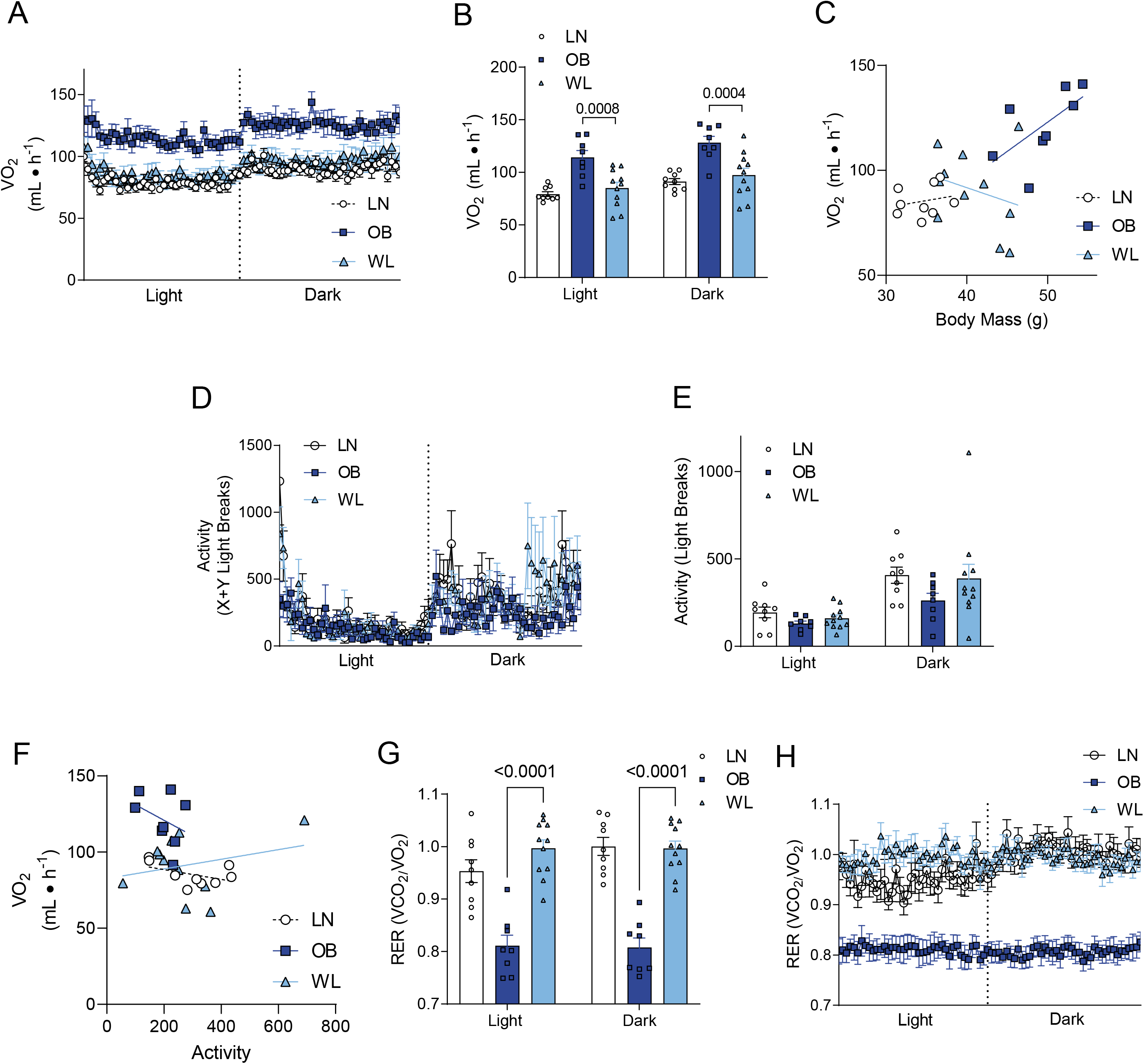
(A&B) Total VO_2_ (unnormalized). (C) Relationship between body mass and total VO_2_ (unnormalized). (D&E) Spontaneous movement. (F) Relationship between spontaneous movement to VO_2_. (G&H) Respiratory exchange ratio (RER). *n*=9 for LN, *n*=8 for OB, *n*=11 for WL. All data are represented as mean ± SEM. Two-way ANOVA with Sidak multiple comparisons (B). p-values indicate statistical significance between OB and WL groups.

**Supplemental Figure S2.**
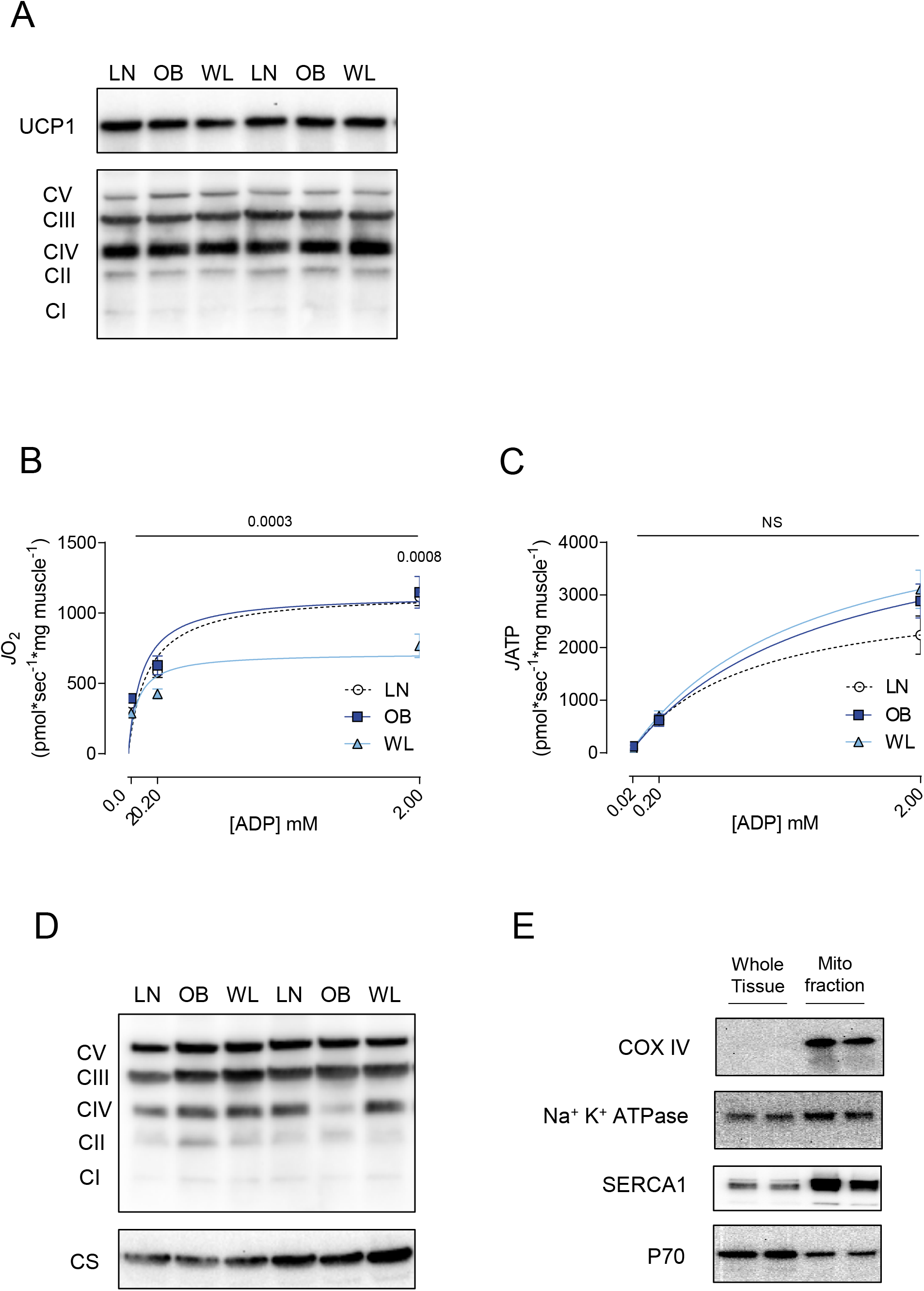
(A) Representative western blots for OXPHOS subunits and UCP1 in homogenates from brown adipose tissues. (B) Rates for oxygen consumption (*J*O_2_). P-value indicates statistical difference between OB and WL groups. (C) Rates for ATP production (*J*ATP). These numbers were used to derive Figure 2C. (D) Representative western blots for OXPHOS subunits and citrate synthase (CS) in total muscle homogenates. (E) Validation of mitochondrial enrichment in the mitochondrial fraction. Percoll gradient isolation was not performed to preserve mitochondrial function for high-resolution respirometry and fluorometry. Blotting for mitochondrial COXIV shows that even a short exposure produces strong immunoreactivity in isolated mitochondrial fraction when such band is barely visible in whole lysate with equal protein loading. Thus, the mitochondrial fraction is highly enriched in mitochondria. In contrast, the mitochondrial fraction also includes proteins from plasma membrane (Na^+^/K^+^-ATPase), endoplasmic reticulum (SERCA), and cytosol (p70) with comparable abundance to whole lysate. Comparable enrichment of total lysate and mitochondrial prep for these organelles is likely due to myofibrillar and extracellular matrix proteins that are highly abundant in whole lysate, making it relatively dilute with intracellular organelles per μg of protein. All data are represented as mean ± SEM. Two-way ANOVA with Sidak multiple comparisons (A,B). p-values indicate statistical significance between OB and WL groups.

**Supplemental Figure S3.**
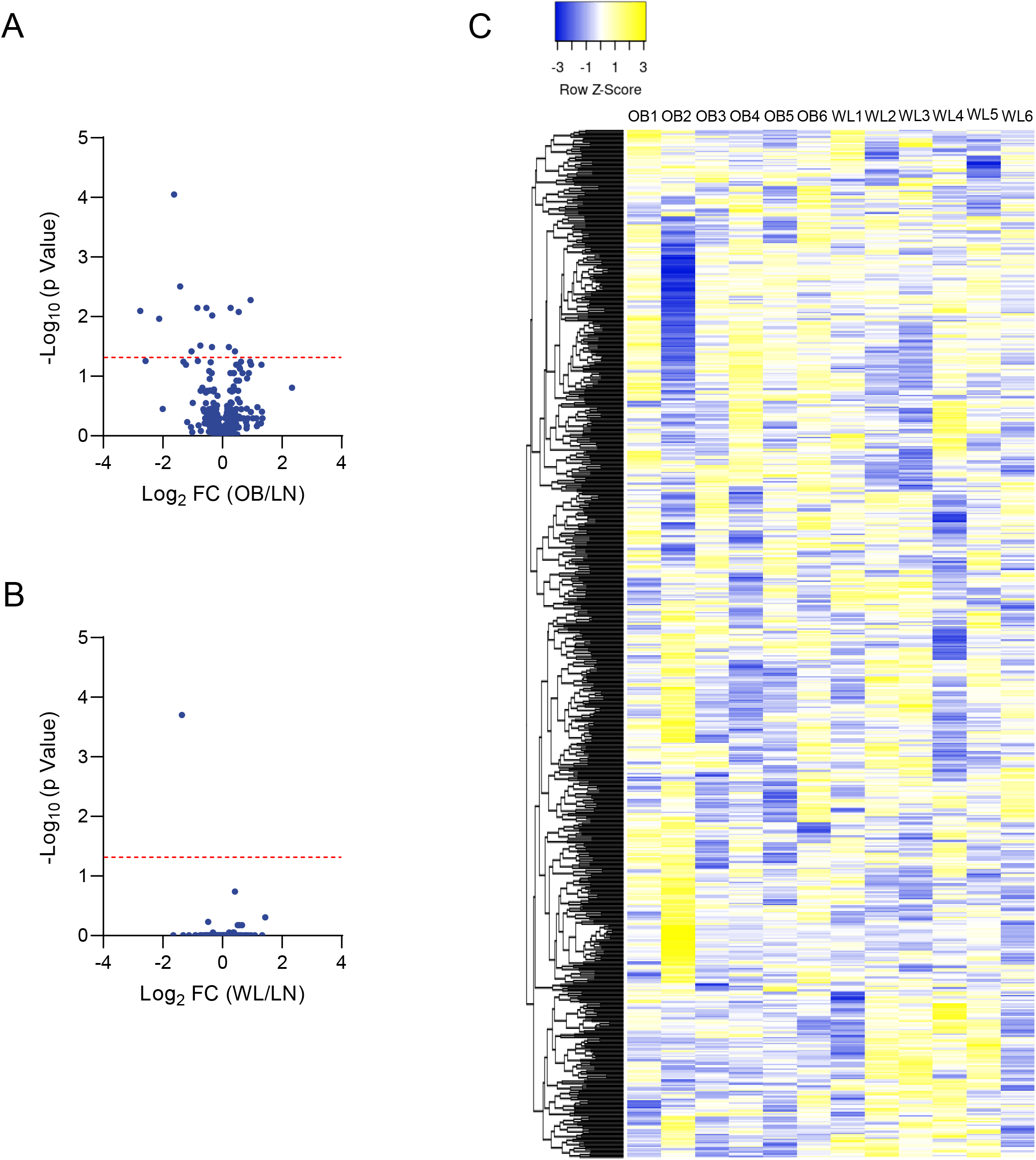
(A) Volcano plot of differentially abundant mitochondria proteins between LN and OB groups. (B) Volcano plot of differentially abundant mitochondria proteins between LN and WL groups. Significance threshold shown with dotted red line for both A and B. (C) Unsupervised clustering analyses of mitochondrial proteome between OB and WL groups.

**Supplemental Figure S4.**
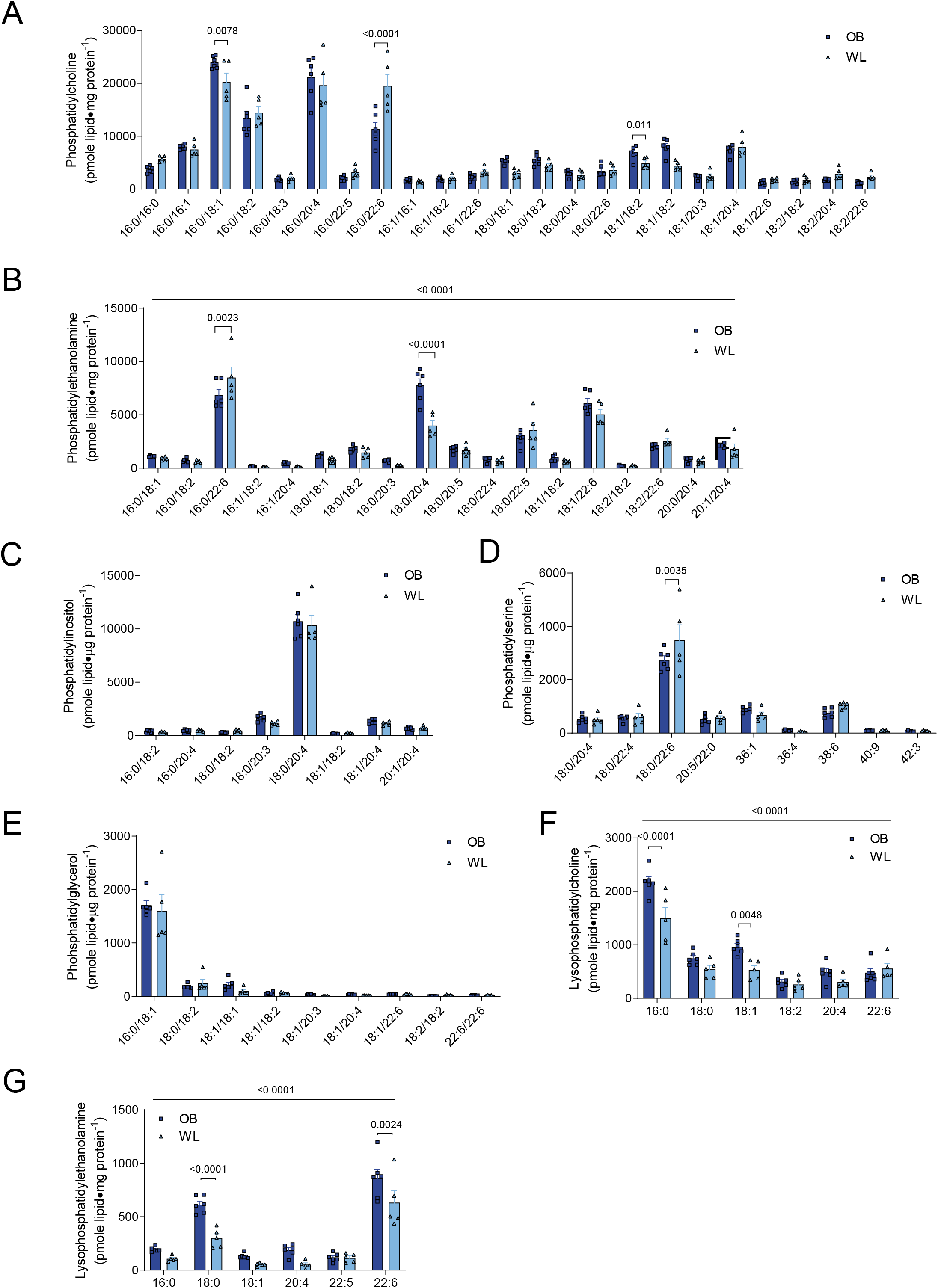
Skeletal muscle mitochondrial lipidomic analyses. (A) Phosphatidylcholine. (B) Phosphatidylethanolamine. (C) Phosphatidylinositol. (D) Phosphatidylserine. (E) Phosphatidylglycerol. (F) Lysophosphatidylcholine. (G) Lysophosphatidylethanolamine. *n*=6 for OB, *n*=5 for WL. All data are represented as mean ± SEM. Two-way ANOVA with Sidak multiple comparisons.

**Supplemental Figure S5.**
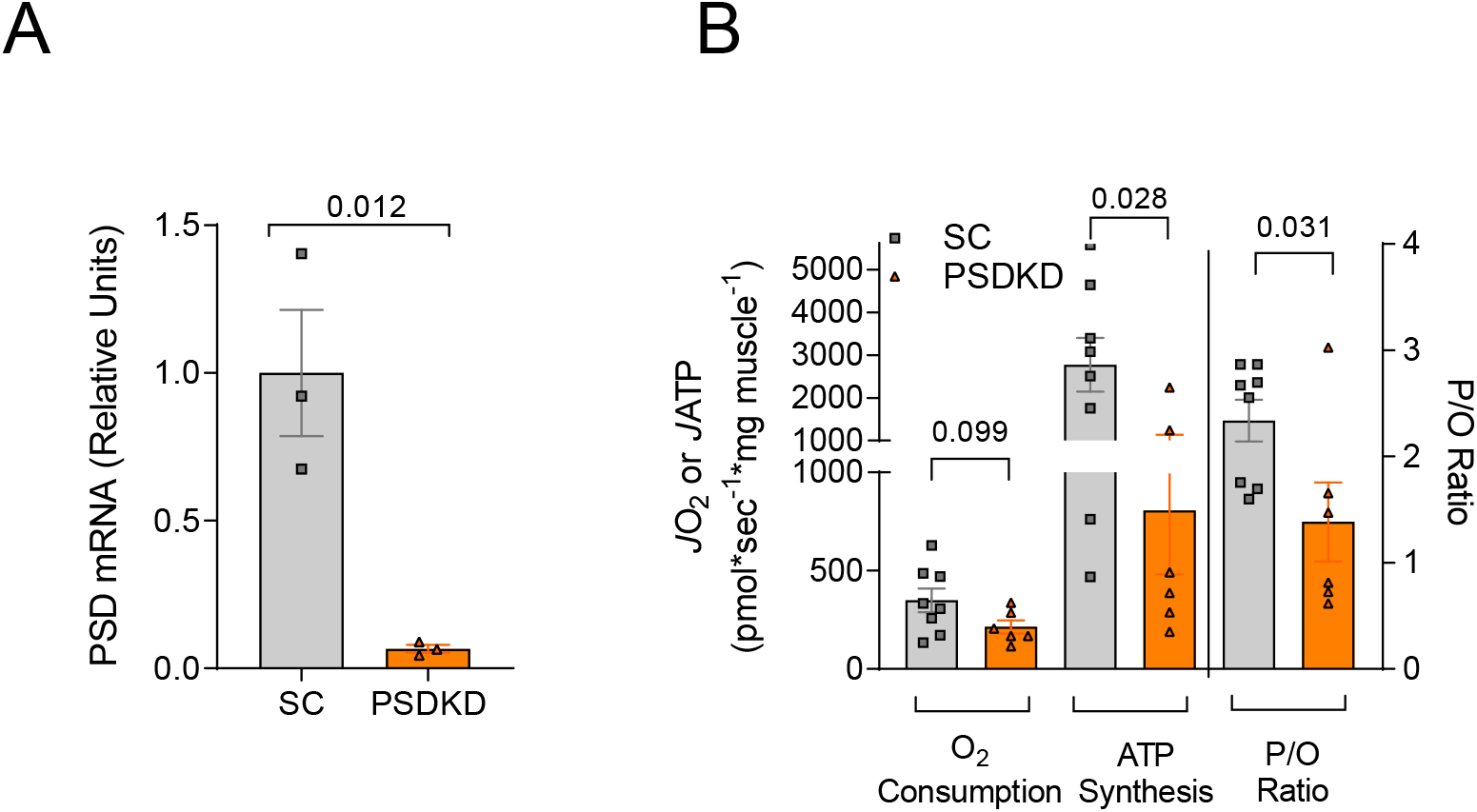
(A) PSD mRNA levels from C2C12 myotubes treated with scrambled (SC) or shPSD (PSDKD). (B) Rates for O_2_ consumption, ATP production, and P/O ratio in isolated mitochondria from SC or PSDKD C2C12 myotubes with 200 μM of ADP (*n*=8 for SC, *n*=6 for PSDKD). All data are represented as mean ± SEM. Unpaired t-test (A,B).

